# Protein profiling of breast carcinomas reveals expression of immune-suppressive factors and signatures relevant to patient outcome

**DOI:** 10.1101/2022.06.03.494654

**Authors:** Felix Ruoff, Nicolas Kersten, Nicole Anderle, Sandra Jerbi, Aaron Stahl, André Koch, Anette Staebler, Andreas Hartkopf, Sara Y. Brucker, Markus Hahn, Katja Schenke-Layland, Christian Schmees, Markus F. Templin

## Abstract

**Background:** In cancerous tissue, a complex interplay of tumour cells with different cell types from the tumour microenvironment is causing modulations in signalling processes. By directly assessing expression of a multitude of proteins and protein variants, extensive information on signalling pathways, their activation status and the effect of the immunological landscape can be obtained providing viable information for treatment response.

**Methods:** Protein extracted from archived breast cancer tissue from patients without adjuvant therapy was subjected to high-throughput Western blotting using the DigiWest technology. Expression of 150 proteins and protein variants covering cell cycle control, apoptosis, Jak/Stat, MAPK-, Pi3K/Akt-, Wnt-, and, autophagic signalling as well as general tumour markers was monitored in a cohort of 84 patient samples. The degree of immune cell infiltration was investigated and set against treatment outcome by integrating patient specific follow-up data.

**Results:** Characterization of the tumour microenvironment by monitoring CD8α, CD11c, CD16 and CD68 expression revealed a strong correlation of event-free patient survival with immune cell infiltration. Furthermore, the presence of tumour infiltrating lymphocytes was linked to a pronounced activation of the Jak/Stat signalling pathway and apoptotic processes. Elevated phosphorylation of peroxisome proliferator-activated receptor gamma (PPARγ, pS112) in non-immune infiltrated tumour tissue suggests a novel immune evasion mechanism in breast cancer characterized by increased PPARγ activation.

**Conclusion:** Multiplexed immune cell marker assessment and protein profiling of tumour tissue provides functional signalling data facilitating breast cancer patient stratification.

## Introduction

Breast cancer is the fifth most common cause of cancer-related deaths worldwide with 2.2 million new cases and around 685.000 deaths in 2020 [1]. A variety of therapeutic options including surgery, chemo-, hormone-, and biological therapy are available and survival rates have increased substantially over the years [2]. Nevertheless, current cancer therapies are lacking a more personalized approach and long-term therapy resistance has become a focus of current research [3, 4]. Novel insights on the establishment, interaction, and control of the tumour immune microenvironment (TIME) have drawn a more comprehensive picture of factors that might account for treatment failure or severe side effects [3, 5]. The detection of tumour infiltrating lymphocytes (TIL) is seen as a major prognostic factor in different carcinomas and has been suggested as a routine pathological evaluation [6, 7]. Large investigations on bulk gene expression of immune infiltrating cells have been performed, associating different immune cell types and risk of relapse [8, 9]. Higher infiltration with immune-cells, is linked to patient outcome and treatment response in several studies [10-13]. Yet, most of these studies focus on the evaluation of single immune cell markers or solely investigate the degree of TIL infiltration most commonly using imaging methods (H&E/IHC/IF staining) for immune cell assessment and subsequent scoring [14, 15]. In the present study, we established the multiplexed assessment of several immune cell markers by DigiWest [16], a multiplexed bead-based western blot, in fresh-frozen tissue samples.

Semi-quantitative protein expression analysis of primary breast cancer tissue allowed for the detection of infiltrating immune cells and a concomitant monitoring of the activation state of central signalling pathways. Activation of immune cell signalling and induction of apoptotic processes was observed in highly immune cell infiltrated tumour tissue, whereas activation of the PPAR gamma signalling was found in tumours tumours with low-level immune cell infiltration.

## Materials and Methods

### Patient Cohort

A retrospective cohort of primary unilateral invasive mamma carcinomas from patients that underwent a primary resection was utilized (snap frozen, n=159, tumour bank University Hospital Tuebingen). Inclusion criteria were hormone receptor and/or human epidermal growth factor receptor (Her2) negative or positive carcinomas, determined by immunohistochemistry at the Institute of Pathology, University of Tuebingen, Germany at the time of surgery. Samples were further classified by occurrence of distant metastases or local relapse within 10 years (poor responder) versus no occurrence of distant metastases, a local relapse within 10 years or contralateral carcinoma within 5 years of primary diagnosis (good responder). In general, exclusion criteria were occurrence of contralateral mamma carcinomas before occurrence of distant metastases in the poor responder-subgroup, respectively within 5 years in the good responder-subgroup as well as presence of a bilateral mamma carcinomas or other malignancies (Figure S1). All patients enrolled neither received any neo-adjuvant treatment nor had any known metastases before surgery.

### Sample preparation and assessment of tumour content

Layered cuts of each fresh-frozen sample were prepared and Hematoxylin-Eosin (H&E) staining was performed according to standard protocols for the first and second layer. Between layers, 100 µm of tissue was trimmed, collected and stored at –80 °C. Prior to protein expression analysis, the prepared layered cuts were H&E stained (see also Figure 1A) and sections 100 µm apart were re-assessed by a pathologist (A.S). The evaluation of these sections revealed that 2.5% (n=4) of the samples were normal tissue, 5.0% (n=8) ductal carcinoma in situ (DCIS) and 1.9% (n=3) mostly contained necrotic tissue. 90.6% (n=144) were classified as invasive ductal carcinoma (IDC) (see also Figure 1B). Yet, 2.5% (n=4) showed a tumour content of app. 5-10%, and 27.7% (n=44) a tumour content in between 15% and 45%. Of all samples, 60.4% (n=96) showed a tumour content of 50% or higher (up to 95%) (see also Figure 1B). Samples with ≥ 50% tumour content were selected for further analysis (n=84) and the intermediate sections from the generated layered cuts were used for protein preparation. In addition, n=10 samples classified as normal tissue or with low tumour content (>10%) were assigned to the baseline control group. Samples were lysed by incubation of collected tissue at 95°C for 10 minutes in lysis buffer (4% LDS, 50mM DTT) (Figure S2).

**Figure 1.**
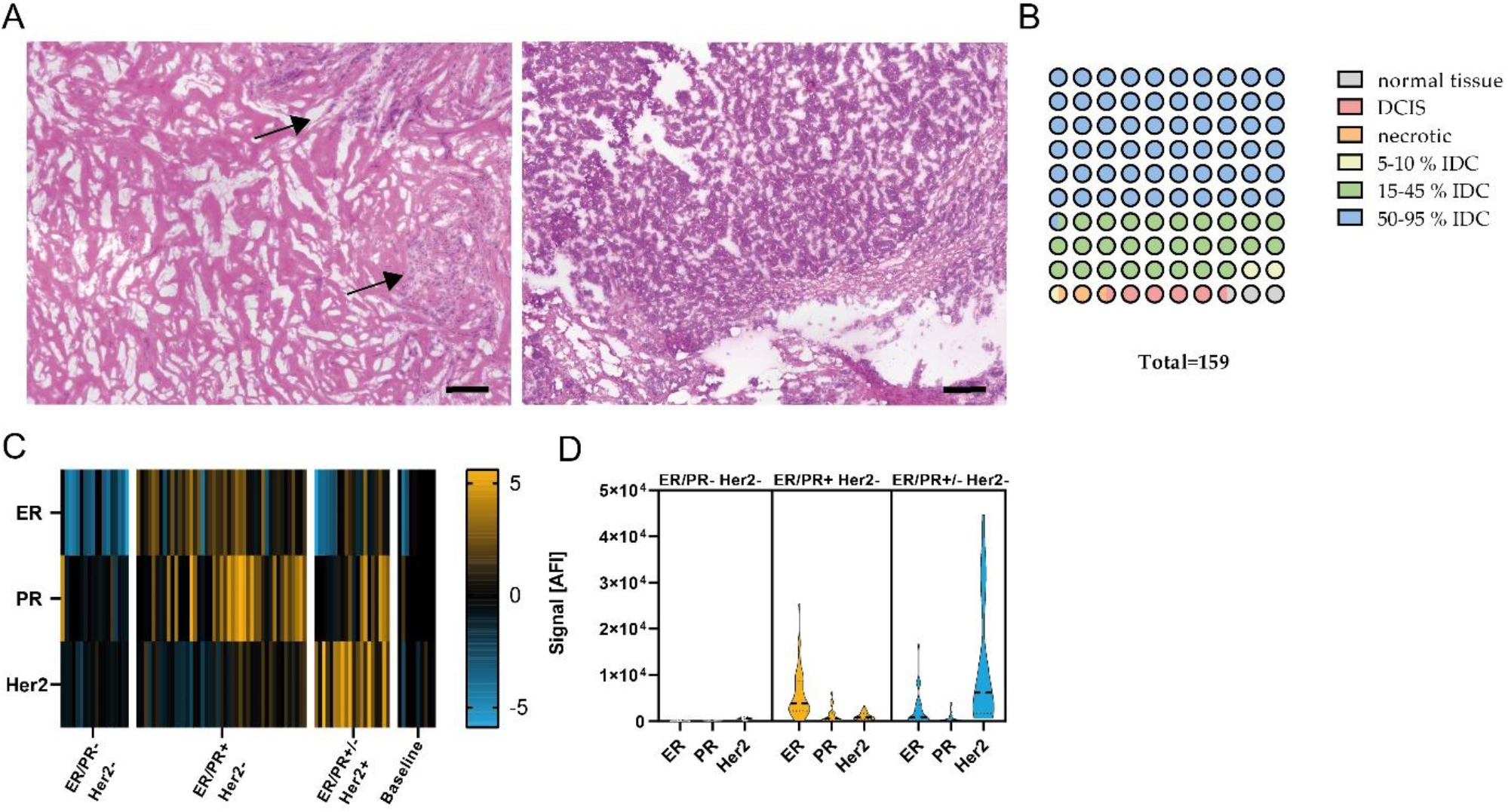
Pathological examination and quality assessment of selected samples. **(A)** Representative images of H&E stained breast carcinomas. **(Left)** app. 20% tumor content. **(Right)** app. 90% tumorcontent. Scale bar, 200 µm. Black arrows, tumor area. **(B)** Pathological classification of entire sample set (1 circle = 1%). n=159. **(C)** Heatmap showing ER, PR and Her2 protein expression in ER/PR-Her2- (n=18), ER/PR+ Her2- (n=45), ER/PR+/- Her2+ (n=20) and Baseline (n=10) subgroup, data median-centered and log2 transformed. Yellow indicates higher expression; blue indicates lower protein expression. **(D)** Violin plots of ER, PR and Her2 expression in ER/PR- Her2-, ER/PR+ Her2- and ER/PR+/-Her2+ subgroups.

To confirm tumour content of the enrolled sample set abundance of the proliferation marker Ki-67 as well as general carcinoma markers Cytokeratin 8/18, Cytokeratin 8 (pS23) and Cytokeratin 6 was assessed by DigiWest analysis (see below). Expression of these markers was found to be significantly different between the tumour sample set and baseline sample sets, revealing a high tumour content in the analysis former sample set (Figure S3A).

### Compliance of receptor status

To review compliance of prepared samples with the pathological evaluation, expression levels of hormone receptors estrogen receptor (ER), progesterone receptor (PR) and human epidermal growth factor receptor (Her2) were performed and resulting signals were compared to pathological receptor status (Figure S3B). A significant difference (P<0.05, Mann-Whintey-*U* test) in signal was found between samples pathologically classified as receptor-positive and receptor-negative or in the baseline group. Samples were categorized into three groups by referencing the pathological receptor status. In ER/PR-Her2-samples (n=18) low or no expression of ER/PR or Her2 was observed. Analysis of ER/PR+ Her2-samples (n=45) showed increased expression of ER and a slight increase in expression in PR but not in Her2. Whereas ER/PR +/-Her2+ samples (n=20) displayed a significant increase in Her2 expression compared to the other groups (see also Figure 1C, D; P<0.05, Mann-Whintey-*U* test). We concluded that DigiWest measurement of hormone receptors and Her2 expression is comparable to the classical pathological assessment of receptor status in the present cohort.

### Immunohistochemical staining

Immunohistochemical staining was performed on 5 µm formalin-fixed paraffin-embedded sections. After de-paraffinization, epitope retrieval was performed at 95°C for 20 minutes in the appropriate antigen retrieval buffer. BLOXXALL-Blocking solution (Vector Laboratories, Burlingame, CA, USA) was added for 10 minutes. After washing in PBS, the sections were incubated with blocking buffer (PBS, 0.25% Triton-X-100, 10% goat serum, 4 drops/ml streptavidin (Vector Laboratories)). Primary antibody diluted in dilution buffer (PBS, 1%BSA, Biotin (Vector Laboratories)) was added and incubated in a humidified chamber. Rabbit (rb) anti-CD8α (#85336, clone D8A8Y, Cell Signaling Technology (CST), 1:100 dilution), rb anit-CD11c (#45581, clone D3V1E, CST, 1:400 dilution), rb anti-CD68 (#76437, MultiMab, CST, 1:200 dilution) and rb anti-CD16 (ab24622, clone EPR14336, Abcam, 1:400 dilution) antibodies were used for staining. Appropriate biotin-conjugated secondary antibody (Jackson Immuno Research, Cambridge, UK) diluted in PBS/1%BSA was added for 30 minutes. After subsequent washing in PBST, slides were incubated with streptavidin-labelled horseradish peroxidase. Peroxidase activity was developed with Novolink 3,3′-Diaminobenzidine (Leicabiosystems, Nußloch, GER). Slides were counterstained with Hematoxylin QS (Vector Laboratories).

Staining for PR, PPARγ and PPARγ—pS112 was performed using a DAKO Autostainer Link 48 (Dako, Jena, GER) and antigen retrieval was performed using a DAKO PT Link (Dako) according to the manufacturer’s recommendations. For staining with mouse anti-PR antibody (IR06861, clone PgR636, Dako) slides were incubated in FLEX TRS HIGH pH buffer (K8004, Dako) at 85 °C for 20 minutes, followed by primary antibody incubation for 20 minutes and incubation with mouse linker for 30 minutes. Subsequently slides were incubated with universal secondary antibody (EnVision FLEX/HRP, K8000, Dako) for 15 minutes. For staining with rb anti-PPARγ (#2435, clone C26H12, CST, dilution 1:100) and rb anti-PPARγ—pS112 (orb5574, Biorbyt, dilution 1:400) slides were incubated in FLEX TRS LOW pH buffer (K8005, Dako) at 85 °C for 20 minutes, followed by primary antibody incubation for 30 minutes and universal secondary antibody for 20 minutes. For detection the EnVision detection system (K500711-2, Dako) was used.

Whole slide images were taken utilizing an Axio Scan Z.1 (Zeiss, Oberkochen, GER). For evaluation of staining intensity, five representative sections of each slide were used and the mean intensity of DAB staining in positive pixel^2^ was calculated utilizing the ZenBlue software 3.1 (Zeiss).

### Multiplex protein profiling via DigiWest

DigiWest was performed as described previously [16]. Briefly, the NuPAGE system (Life Technologies, Carlsbad, USA) with a 4-12% Bis-Tris gel was used for gel electrophoresis and Western blotting onto PVDF membranes. After washing with PBST, proteins were biotinylated by adding 50 µM NHS-PEG12-Biotin in PBST for 1 h to the Membrane. After washing in PBST Membranes were dried overnight. Each Western-Blot lane was cut into 96 strips of 0.5 mm each. Strips of one Western blot lane were sorted into a 96-Well plate (Greiner Bio-One, Frickenhausen, GER) according to their molecular weight. Protein elution was performed using 10 µl of elution buffer (8 M urea, 1% Triton-X100 in 100 mM Tris-HCl pH 9.5). Neutravidin coated MagPlex beads (Luminex, Austin, TX, USA) of a distinct color ID were added to the proteins of a distinct molecular weight fraction and coupling was performed overnight. Leftover binding sites were blocked by adding 500 µM deactivated NHS-PEG12-Biotin for 1 h. To reconstruct the original Western blot lane, the beads were pooled, at which the color IDs represent the molecular weight fraction of the proteins.

For antibody incubation 5 µl of the DigiWest Bead mixes were added to 50 µl assay buffer (Blocking Reagent for ELISA (Roche, Rotkreuz, Switzerland) supplemented with 0.2% milk powder, 0.05% Tween-20 and 0.02% sodium azide) in a 96 Well plate. Assay buffer was discarded and 30 µl of primary antibody diluted in assay buffer was added per well. Primary antibodies were incubated overnight at 15 °C on a shaker. Subsequently, were washed twice with PBST. After washing, 30 µl of species specific-secondary antibody diluted in assay buffer labeled with phycoerythrin was added and incubation took place for 1 h at 23°C. Before the readout on a Luminex FlexMAP 3D instrument beads were washed twice with PBST.

Analysis and Peak integration were performed by utilizing the novel DigiWest-Analyzer software package [17].

### Statistical analysis

Statistical comparison was done by using Mann-Whitney-*U* test. (GraphPad Prism version 9.2.0, GraphPad Software). Spearman correlation analysis, Hierarchical cluster analysis, Chi-square test, Kaplan-Meier plot and log-rank test were carried out utilizing the DigiWest-Evaluator software package [17]. P values of <0.05 were considered statistically significant if not stated differently.

### Pathway enrichment analysis

Testing for significantly enriched pathways was performed with an over representation analysis using Fisher’s exact test with subsequent calculation of Storey’s Q-values for multiple testing correction. The subsets of analytes that were used for this analysis were defined by applying the Mann-Whitney-*U* test to identify differentially expressed analytes between the good responder and poor responder groups. The pathway enrichment pipeline was carried out utilizing the DigiWest-Evaluator software package [17].

## Results

### Sample quality control and DigiWest protein expression analysis

After initial sample assessment (n=159, Figure 1A, B) samples with tumour content >50% and sufficient protein amount (n=84) as well as control samples with 10% or less tumour content (n=10), were selected for extensive protein expression analysis. To identify markers relevant for the differentiation of good and poor responders we measured 150 proteins and protein variants using DigiWest, mainly focusing on functional signal transduction i.e. protein phosphorylation (covering 41 phospho-variants). This extensive expression analysis encompassed cell cycle control, apoptosis, Jak/Stat-, MAPK-, Pi3K/Akt-, Wnt- and autophagic signalling pathways as well as general tumour and immune-cell markers. DigiWest evaluation of hormone receptor and Her2 receptor expression complied with the pathologically assessed receptor status, confirming high quality of the selected tumour samples (Figure 1C, D).

To examine the connection of cellular signalling and responder status, PANTHER pathway enrichment analysis was conducted for all proteins differentially expressed between good and poor responder samples [18, 19]. The highest –log2 Q-value was found for the Jak/Stat signalling pathway (Figure 2A). Additionally, members of Jak/Stat signalling, and several immune cell markers displayed a significant difference when comparing protein expression of good and poor responder samples (Mann-Whitney-*U* test, FDR limit 0.1, Figure 2B). Taken together these results indicate a connection of immune-cell related signalling pathway activity and patient treatment response.

**Figure 2.**
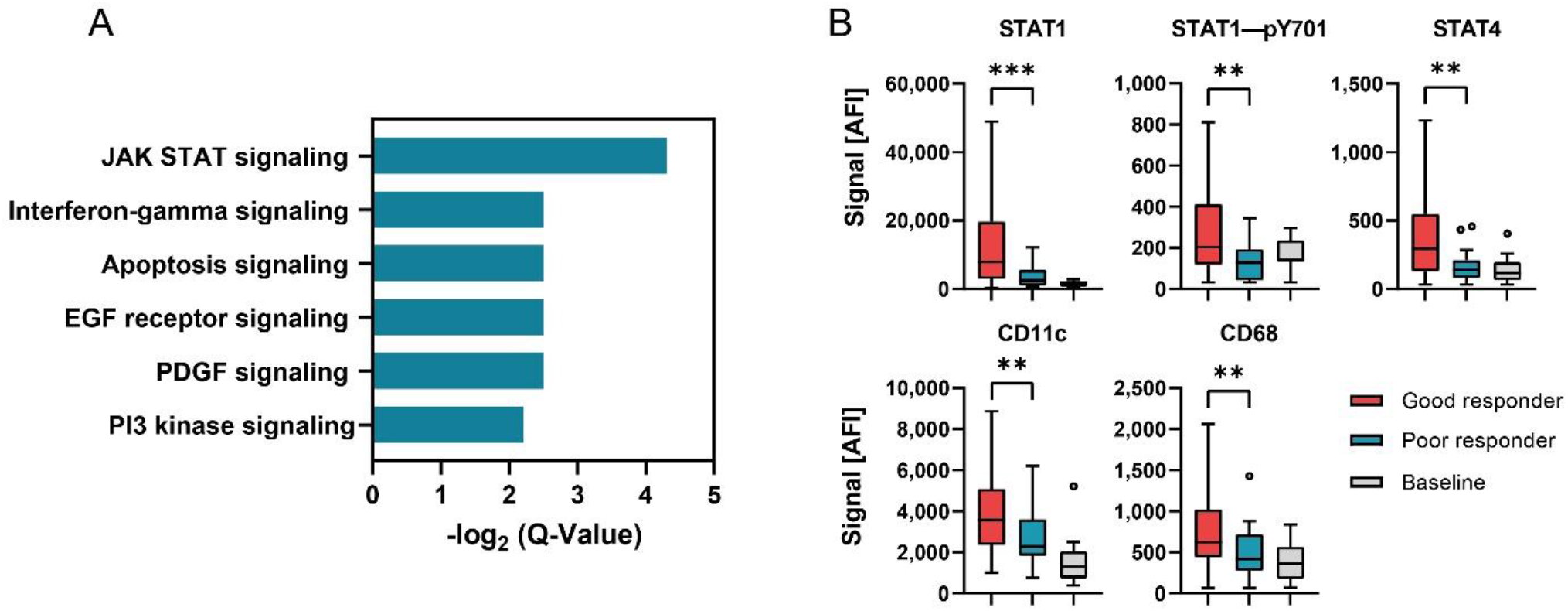
Responder status depended differences in signal transduction. **(A)** –log2 Q-Values of PANTHER pathway analysis as bar graphs. **(B)** Protein expression of selected analytes displaying significant differences in expression between responder groups. Data shown as box-whisker-plots for good responder (n=58), poor responder (n=21) and baseline (n=10) subgroup. ***P<0.001, **P<0.01. Mann-Whitney-*U* test.

#### Patient stratification based on immune cell infiltration analysis by DigiWest

The degree and type of tumour infiltration by immune cells has become a novel and promising stratification factor for evaluating patient outcome. Therefore, we evaluated the subset of measured immune cell markers in more detail. Correlation analysis of CD8α, CD4, CD68, CD11c, CD16, CD56, CD25 as well as CD163 protein expression was performed. CD8α, CD68, CD11c and CD16 displayed the highest correlation (Spearman’s r < 0.55, Figure 3A, Table S1), suggesting a co-occurrence of represented immune cells.

**Figure 3.**
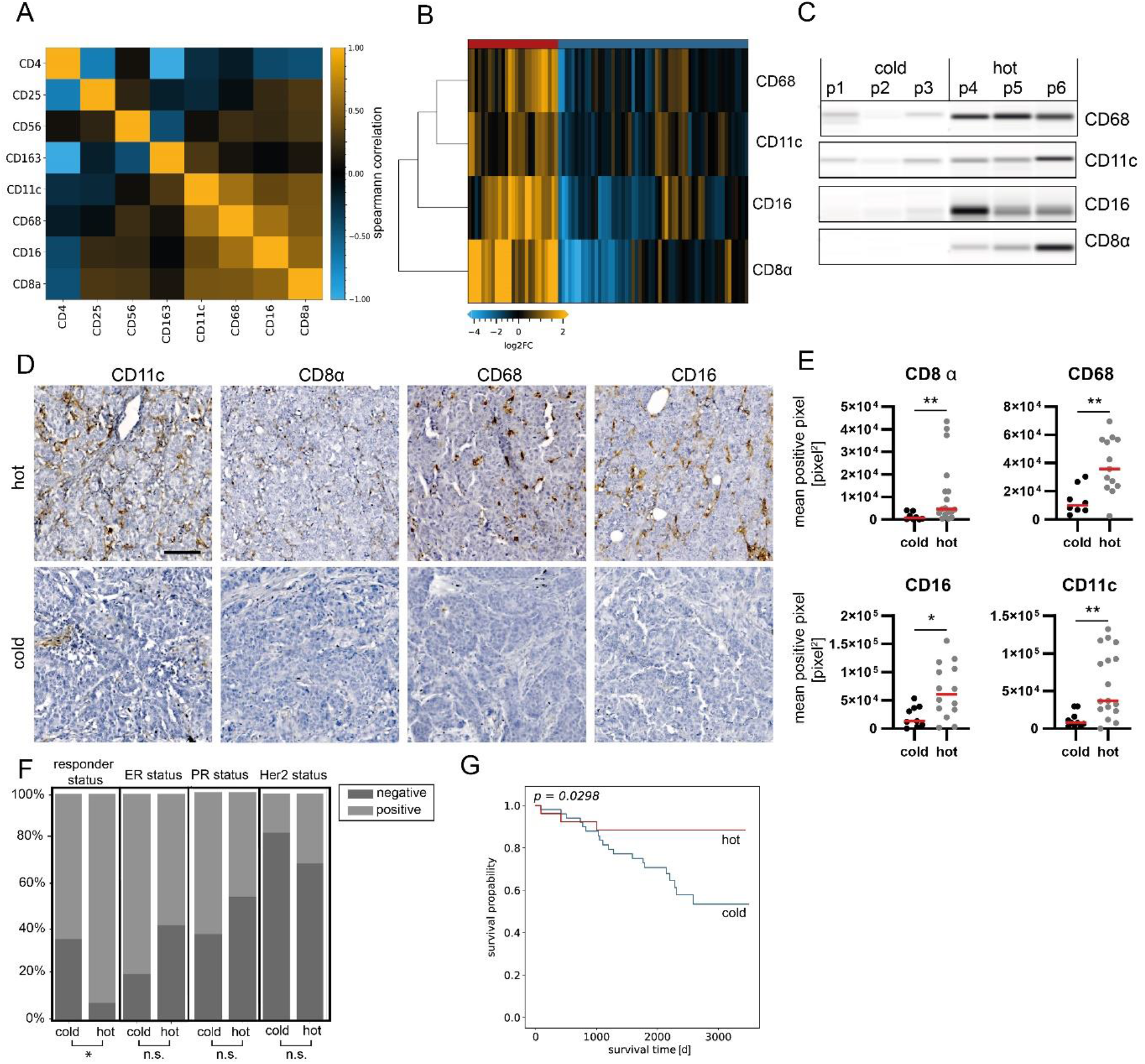
Sample stratification based on immune marker assessment. **(A)** Correlation plot (Spearman’s correlation) of immune cell marker expression in the analysis sample set. Highest correlation was found for CD8α, CD11c, CD68 and CD16 (r > 0.55). n=84. **(B)** Heatmap showing protein expression levels of CD8α, CD11c, CD68 and CD16. Hierachical clustering of analytes and samples with Euclidean distance and complete linkage. Data are normalized to total protein, centered on median of all samples and log2 transformed. **(C)** Representative western-blot mimics of CD8α, CD11c, CD68 and CD16 (grayscale maps generated from the DigiWest data). For graphical representation, background-subtracted raw data from representative hot and cold samples were used. n=3. **(D)** Representative images of CD11c, CD8, CD68 and CD16 immunohistochemical staining in hot and cold samples. Scale bar, 50 µm. **(E)** Mean positive pixel values of 5 representative 10x sections per available FFPE sample for hot and cold carcinomas classified by DigiWest. CD8α: cold n=8, hot n=17; CD68: cold n=7, hot n=13; CD16: cold=9, hot n=14; CD11c: cold n=10, hot n=17. Mann-Witney-*U* test, **P<0.01; *P<0.05. **(F)** Distributions for responder, ER, PR and Her2 status as percentages stratified by infiltration status. Chi-square-test, *P<0.05, ns indicates no significant difference. **(G)** Kaplan-Meier analyses of event-free survival in patients stratified by infiltration status. P=0.0298, Log-rank test.

Unsupervised hierarchical cluster linkage analysis of CD8α (common marker for cytotoxic t-killer cells), CD16 (common marker for cytotoxic natural killer (NK) cells), CD11c (marker for dendritic cells) and CD68 (general marker for macrophages) revealed two distinct sample groups with different levels of immune marker expression (Euclidean distance, complete linkage, Figure 3B, C). The group with higher immune cell marker expression (n=27) will be referred to as “hot tumours”, whereas the group with lower immune cell marker expression (n=57) will be referred to as “cold tumours”.

Subsequently, CD8α, CD68, CD11c and CD16 were immunohistochemically stained on matched FFPE sections, when available (Figure 3D, Figure S4). Concomitantly, a significant difference in mean pixel intensity among hot and cold tumour samples categorized by DigiWest was detected (Figure 3E; Mann-Whitney-*U* test, P<0.05).

When comparing various clinical variables such as the type of surgery, age and most notably receptor status (ER, PR or Her2) we did not observe any difference between hot and cold breast carcinomas. Importantly, the responder status was the only clinical variable significantly enriched within the hot tumour group (Figure 3F; Table 1; P<0.05, chi-square test).

**Table 1.** Patient and tumour characteristics, of all samples included in the analysis sample set, stratified by responder and immune infiltration status as indicated by DigiWest. P: Chi Square or Wilcoxon rank-sum test between the two groups.

| Characteristic | Overall (N=84) |  | Poor responder (N=21) |  | Good responder (N=58) |  | P | Cold (N=57) |  | Hot (N=27) |  | P |
| --- | --- | --- | --- | --- | --- | --- | --- | --- | --- | --- | --- | --- |
|  | No. of Patients | % | No. of Patients | % | No. of Patients | % |  | No. of Patients | % | No. of Patients | % |  |
| **Follow-up, years** |  |  |  |  |  |  | 0.49 |  |  |  |  | 0.7 |
| Median (range) | 6.1 (0.2-9.9) |  | 6.3 (0.2-9.9) |  | 6.2 (0.3-9.6) |  |  | 6.3 (0.2-9.9) |  | 6.2 (0.9-9.4) |  |  |
| **Age at surgery, years** |  |  |  |  |  |  | 0.13 |  |  |  |  | 0.4 |
| Median (range) | 61 (30-85) |  | 66 (41-85) |  | 58 (84-30) |  |  | 60 (31-85) |  | 62 (30-84) |  |  |
| **Tumor size (cm)** |  |  |  |  |  |  | 0.15 |  |  |  |  | 0.3 |
| <2 | 23 | 27.4 | 4 | 19.0 | 17 | 29.3 |  | 14 | 24.6 | 8 | 29.6 |  |
| 2-5 | 55 | 65.5 | 13 | 61.9 | 39 | 67.2 |  | 37 | 64.9 | 19 | 70.4 |  |
| >5 | 6 | 7.1 | 4 | 19.0 | 2 | 3.4 |  | 6 | 10.5 | 0 | 0 |  |
| **Nodal status** |  |  |  |  |  |  | 0.4 |  |  |  |  | 0.3 |
| Negative | 47 | 56.0 | 10 | 47.6 | 35 | 60.34 |  | 29 | 50.9 | 18 | 66.7 |  |
| Positive | 36 | 42.9 | 11 | 52.4 | 22 | 37.93 |  | 27 | 47.4 | 9 | 33.3 |  |
| Unknown | 1 | 1.2 | 0 | 0.0 | 1 | 1.72 |  | 1 | 1.8 | 0 | 0 |  |
| **Hormone receptor status** |  |  |  |  |  |  |  |  |  |  |  |  |
| ER-Positive | 24 | 28.6 | 15 | 71.4 | 42 | 72.4 | 0.84 | 44 | 77.2 | 16 | 59.3 | 0.2 |
| ER-Negative | 60 | 71.4 | 6 | 28.6 | 16 | 27.6 |  | 13 | 22.8 | 11 | 40.7 |  |
| PR-Positive | 38 | 45.2 | 12 | 57.1 | 33 | 56.9 | 0.81 | 34 | 59.6 | 12 | 44.4 | 0.3 |
| PR-Negative | 46 | 54.8 | 9 | 42.9 | 25 | 43.1 |  | 23 | 40.4 | 15 | 55.6 |  |
| **HER2 status** |  |  |  |  |  |  | 0.83 |  |  |  |  | 0.5 |
| Positive | 20 | 23.8 | 4 | 19.0 | 14 | 24.1 |  | 11 | 19.3 | 8 | 29.6 |  |
| Negative | 63 | 75.0 | 17 | 81.0 | 43 | 74.1 |  | 45 | 78.9 | 18 | 66.7 |  |
| Unknown | 1 | 1.2 | 0 | 0.0 | 1 | 1.7 |  | 1 | 1.8 | 1 | 3.7 |  |
| **Type of surgery** |  |  |  |  |  |  | 0.27 |  |  |  |  | 0.4 |
| BCS | 47 | 56.0 | 12 | 57.1 | 34 | 58.6 |  | 34 | 59.6 | 13 | 48.1 |  |
| SSM | 1 | 1.2 | 1 | 4.8 | 0 | 0.0 |  | 1 | 1.8 | 0 | 0 |  |
| Ablatio | 16 | 19.0 | 3 | 14.3 | 10 | 17.2 |  | 5 | 8.8 | 5 | 18.5 |  |
| Mastectomy | 10 | 11.9 | 5 | 23.8 | 5 | 8.6 |  | 2 | 3.5 | 0 | 0 |  |
| Quadrantectomy | 2 | 2.4 | 0 | 0.0 | 1 | 1.7 |  | 10 | 17.5 | 5 | 18.5 |  |
| Segmental resection | 1 | 1.2 | 0 | 0.0 | 1 | 1.7 |  | 0 | 0 | 1 | 3.7 |  |
| Mastopexy | 5 | 6.0 | 0 | 0.0 | 5 | 8.6 |  | 3 | 5.3 | 2 | 7.4 |  |
| NSM | 1 | 1.2 | 0 | 0.0 | 1 | 1.7 |  | 1 | 1.8 | 0 | 0 |  |
| Unknown | 1 | 1.2 | 0 | 0.0 | 1 | 1.7 |  | 1 | 1.8 | 0 | 0 |  |
| **Responder status** |  |  |  |  |  |  |  |  |  |  |  | 0.02 |
| Poor responder | 21 | 25.0 | - | - | - | - |  | 19 | 33.3 | 2 | 7 |  |
| Good responder | 58 | 69.0 | - | - | - | - |  | 34 | 59.6 | 24 | 89 |  |
| Unknown | 5 | 6.0 |  |  |  |  |  | 4 | 7.0 | 1 | 4 |  |

Tumour infiltration with immune cells is generally associated with patient outcome and event-free survival (EFS, time from definitive surgery until disease recurrence/metastases or death from any cause). Therefore, we reviewed the difference in clinical outcome after primary surgery in the present cohort. The group classified as “hot tumours” indeed had a significantly better outcome when evaluating 10 years EFS (Figure 3G; P=0.03, log-rank test). By comparing EFS between hot and cold tumour samples in subgroups categorized through pathological hormone and Her2 receptor status a significant difference was found in ER+ and PR+ samples; yet, more strongly infiltrated samples displayed a tendency towards better EFS (Figure S5). Univariate Cox regression confirmed that immune cell infiltration status was a prognostic factor of clinical outcome (P=0.04).

### Focused protein expression analysis of hot and cold breast carcinomas

Next, we performed a detailed analysis to identify differential protein expression in breast carcinomas classified as hot or cold tumours. We allocated the samples to the hot or cold tumour group by assessment of immune cell markers as described above and compared protein expression levels. N=30 analytes displayed a significant difference in expression (Mann-Whitney-*U* test; FDR Benjamini-Hochberg; corrected P < 0.05) and a log2 fold change of at least 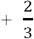 or 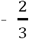 (Figure 4A, B, D; see Table 2 for all significantly differentially expressed proteins; Figure S6).

**Figure 4.**
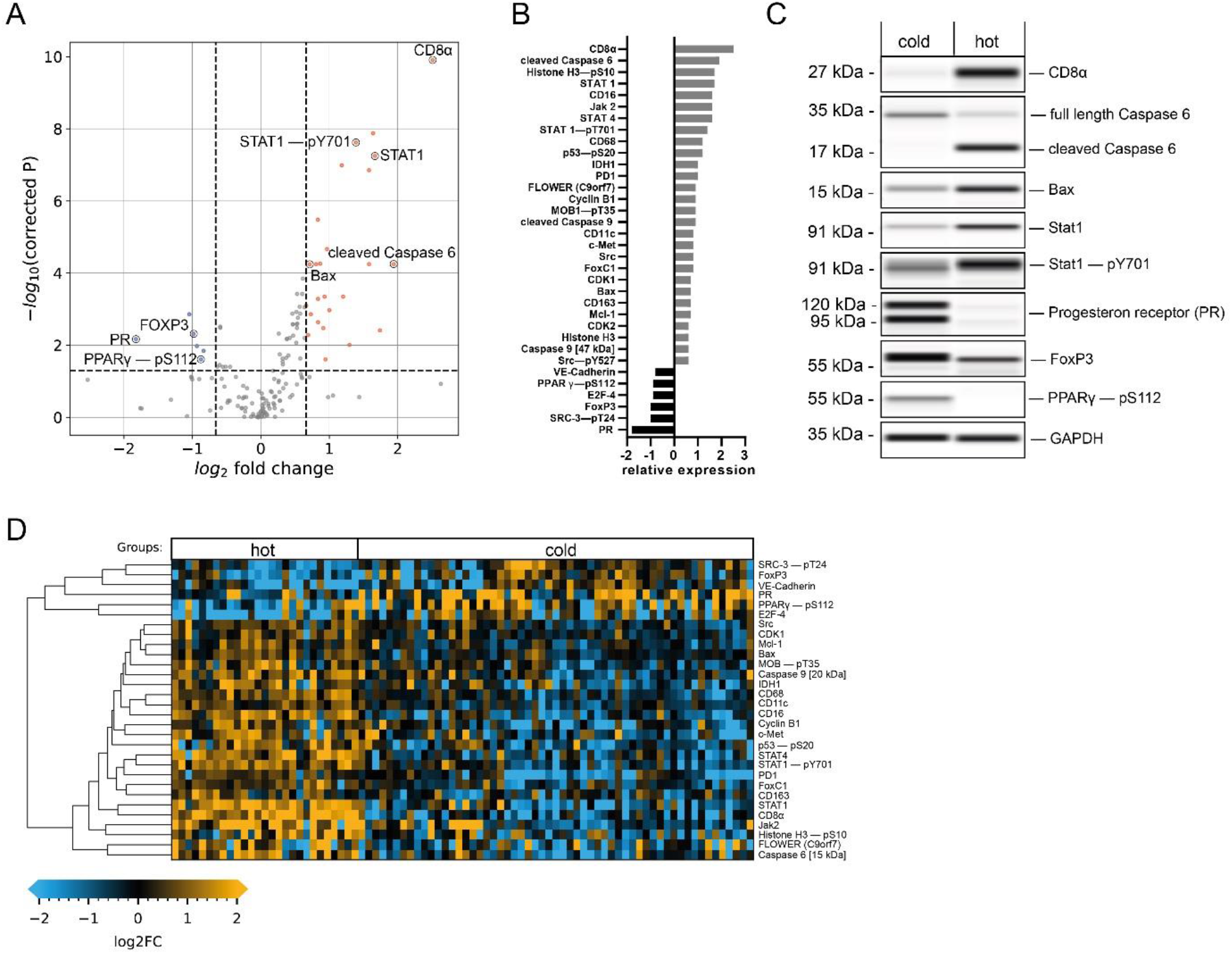
Immune cell infiltration-dependent changes in protein expression. **(A)** Vulcano plot (-log10 corrected P-value versus log2 fold change) depicting differences in protein expression between hot (n=27) and cold (n=57) samples. Data from 150 proteins and protein variants were analysed. Wilcoxon rank sum test with Benjamini-Hochberg multiple testing correction; horizontal dashed line indicates a P*-*value of 0.05; vertical dashed line indicates a log2 fold change of at least 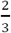. Blue and red dots indicate analytes with at least 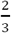 fold difference in median expression and a P-value below 0.05. **(B)** Bar graphs of log2-transformed ratios calculated from mean protein expression of high and low infiltrated samples. Analytes are sorted from the highest positive change to the highest negative change. **(C)** Western-blot mimics of selected analytes from two representative patients (grayscale maps generated from the DigiWest data. For graphical representation, background-subtracted raw data were used. **(D)** Heatmap of analytes with significantly different expression between hot and cold samples displaying a fold change greater than 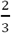. Hierachical clustering of analytes using Euclidean distance and complete linkage.

**Table 2.** Protein analytes displaying significantly different expression between hot and cold breast tumour samples (Mann-Whitney-U test; FDR Benjamini-Hochberg; corrected P < 0.05).

| Analyte | uncorrected *p* value | corrected *p* value | log2 fold |
| --- | --- | --- | --- |
| PR | 0.002 | 0.007 | -1.8 |
| SRC-3—pT24 | < 0.001 | 0.001 | -1 |
| FoxP3 | 0.001 | 0.005 | -1 |
| E2F-4 | 0.003 | 0.011 | -0.9 |
| PPAR gamma—pS112 | 0.009 | 0.025 | -0.9 |
| VE-Cadherin | 0.005 | 0.015 | -0.8 |
| Cytokeratin 8/18 | 0.016 | 0.040 | -0.6 |
| Glycogen Synthase—pS641 | 0.014 | 0.036 | -0.6 |
| PTEN | 0.001 | 0.003 | -0.6 |
| PDK1 | 0.001 | 0.003 | -0.6 |
| PTEN—pS380 | 0.017 | 0.041 | -0.6 |
| mTOR | 0.016 | 0.040 | -0.5 |
| Dvl2 | 0.016 | 0.040 | -0.4 |
| MAD2L1 | 0.001 | 0.005 | 0.3 |
| E2F-1 | 0.006 | 0.018 | 0.3 |
| p70 S6 kinase—pT389 | 0.008 | 0.024 | 0.3 |
| Erk1/2 | 0.011 | 0.030 | 0.3 |
| IKK alpha | 0.013 | 0.035 | 0.3 |
| p38 MAPK—pT180/Y182 | 0.008 | 0.023 | 0.4 |
| PD-L1 | 0.002 | 0.007 | 0.4 |
| A-Raf | 0.002 | 0.007 | 0.4 |
| TSG101 | 0.007 | 0.020 | 0.4 |
| NF-kB p65—pS468 | 0.002 | 0.006 | 0.4 |
| PD-L1 | < 0.001 | 0.001 | 0.5 |
| NF-kB p105/p50 [50 kDa] | 0.001 | 0.003 | 0.5 |
| CD25 | 0.001 | 0.003 | 0.5 |
| c-Raf—pS259 | 0.002 | 0.006 | 0.5 |
| PI3-kinase p85 | 0.003 | 0.009 | 0.5 |
| CD56 | < 0.001 | 0.001 | 0.5 |
| p38 MAPK | 0.001 | 0.003 | 0.5 |
| GSK3 beta | 0.001 | 0.003 | 0.5 |
| RUNX2 | < 0.001 | 0.000 | 0.5 |
| p53—pS37 | 0.001 | 0.003 | 0.5 |
| STAT 5 alpha | < 0.001 | 0.002 | 0.5 |
| MEK2 | < 0.001 | 0.001 | 0.6 |
| Caspase 3 | < 0.001 | 0.001 | 0.6 |
| Src—pY527 | 0.008 | 0.024 | 0.6 |
| Caspase 9 [47 kDa] | < 0.001 | < 0.001 | 0.6 |
| Histone H3 | < 0.001 | < 0.001 | 0.6 |
| CDK2 | 0.002 | 0.007 | 0.6 |
| Mcl-1 | < 0.001 | 0.001 | 0.7 |
| CD163 | 0.001 | 0.005 | 0.7 |
| Bax | < 0.001 | < 0.001 | 0.7 |
| CDK1 | < 0.001 | 0.001 | 0.7 |
| FoxC1 | < 0.001 | < 0.001 | 0.8 |
| Src | < 0.001 | 0.001 | 0.8 |
| c-Met | < 0.001 | 0.002 | 0.8 |
| CD11c | < 0.001 | < 0.001 | 0.8 |
| Caspase 9 [35 kDa] | < 0.001 | < 0.001 | 0.9 |
| MOB1—pT35 | 0.001 | 0.003 | 0.9 |
| Cyclin B1 | < 0.001 | < 0.001 | 0.9 |
| FLOWER (C9orf7) | 0.009 | 0.025 | 0.9 |
| PD1 | < 0.001 | < 0.001 | 1 |
| IDH1 | < 0.001 | 0.001 | 1 |
| p53—pS20 | < 0.001 | < 0.001 | 1.2 |
| CD68 | < 0.001 | < 0.001 | 1.2 |
| STAT 1—pT701 | < 0.001 | < 0.001 | 1.4 |
| STAT 4 | < 0.001 | < 0.001 | 1.6 |
| Jak 2 | < 0.001 | < 0.001 | 1.6 |
| CD16 | < 0.001 | < 0.001 | 1.6 |
| STAT 1 | < 0.001 | < 0.001 | 1.7 |
| Histone H3—pS10 | 0.001 | 0.004 | 1.7 |
| Caspase 6 [15 kDa] | < 0.001 | < 0.001 | 1.9 |
| CD8a | < 0.001 | < 0.001 | 2.5 |

Looking into detail our data show that CD163 a marker for M2 type tumour-associated macrophages (TAMs), linked to tumour progression [20], displayed elevated expression levels (0.7-fold increase) in the hot tumour group. Expression of CD56, used for the identification of natural killer cells [21], and IL-2Rα/CD25, characteristic for regulatory t-cells (Tregs) [22] was also increased in this subgroup. Interestingly, the transcription factor Foxp3, which is a common marker for Tregs broadly linked to immunosuppression and tumour protection [23, 24] showed significantly increased expression in the cold tumour subgroup (Figure 4C).

Additionally, increased levels of various members of the Jak/Stat pathway were detected in the hot tumour subgroup. STAT4, a known mediator of IL-12 response [25] as well as STAT1, known to be essential for interferon-α (IFN-α) and IFN-γ response, as well as its active phosphor-variant (Tyr701) [26] were significantly increased in hot tumours (Figure 4C). Furthermore, the Janus tyrosin kinase 2 (Jak2), an important cytokine receptor [27], displayed a 1.6 fold higher elevated expression in hot tumours. The programmed cell death 1 protein (PD-1), which is known to be an important regulator of immune cell activity [28] was also enriched in this group. Thus, higher immunosurveillance in the hot carcinoma group is characterized by increased expression levels of additional immune cell markers and important members for immune relevant signalling pathways supporting the previously established tumour groups.

#### Hot tumour samples show increased proliferative activity and a more competitive phenotype

Expression levels of the proto-oncogene tyrosine kinase Src as well as the calcium channel flower (FLOWER) were found to be increased in hot tumour samples. Src plays a critical role in multiple cellular processes, including proliferation and invasion and can be a driver for uncontrolled cell growth [29]. Expression of FLOWER, a cellular fitness sensor has been associated with a competitive growth advantage of cancer cells [30, 31]. In addition, the hepatocyte growth factor receptor (c-Met), which is instrumental for increased cell growth and associated with aggressive cancer phenotypes [32, 33] was highly expressed in samples with higher immune cell infiltration. Additionally, the transcription factor forkhead box C1 (FoxC1), which is linked to breast cancer invasiveness [34, 35] was highly expressed in samples assigned to the hot tumour subgroup. Moreover, expression of vascular endothelial cadherin (VE-Cadherin), which is important for adhesion of cancer cells to the endothelium and is lost during advanced phases of mesenchymal-to-epithelial transition (MET) [36], was decreased as compared to the cold tumour group. Taken together these results indicate a more competitive phenotype in hot breast tumour tissue samples.

Additionally, a significant promotion of the cell cycle was found in the hot tumour subgroup, as indicated by higher expression of the important regulatory proteins Cyclin B1 and CDK1, responsible for G2/M-phase progression [37]. This was accompanied by an increase of phosphorylation of the mitosis marker Histone H3 at Serine 10 [38], as well as significant decrease in expression of the cell growth repressor E2F-4 [39].

In conclusion, it is conceivable that the higher expression of proliferative and competitive signalling proteins facilitate the immune recognition of breast cancer cells and therefore lead to higher rates of immune cell infiltration.

#### Elevated immune cell infiltration induces expression of tumour suppressive markers and apoptotic activity

Our data also indicate that hot tumour samples show a significant increase in expression of several tumour suppressors. Phosphorylation of the tumour suppressor p53, at Serine 20, which enhances p53 activity leading to cell cycle arrest or apoptosis [40] was detected and a 1.2 fold increase of phosphorylation was observed. In addition, increased expression of isocitrat-dehyndrogenase 1 (IDH1), another suppressor of tumorigenesis [41], and of phosphorylated and activated MOB1 (pT35), a member of the Hippo pathway [42], was observed in hot tumour samples.

Consequently, this subgroup displayed an increase in apoptotic activity, indicated by enriched expression of several pro-apoptotic proteins. These include the initiator cysteine aspartic acid protease 9 (Caspase 9), serving as an amplifier of the apoptotic response [43] and Caspase 6, (one of the major executioner caspases). Both promote apoptosis in its cleaved and active form [44]. Crucially, we observed an upregulation of the cleaved caspase 9 fragment at 35 kDa as well as the cleaved Caspase 6 fragment at 15 kDa (Figure 4C), but not of the full-length proteins in the hot tumour subgroup. Similarly expression of, the pro-apoptotic factor Bax, [45], was significantly enriched in samples with higher immune infiltration. Interestingly, the anti-apoptotic Bcl-2 family member Mcl-1, which antagonizes pro-apoptotic Bcl-2 proteins [46] showed a similar trend. These results suggest that higher immune cell infiltration leads to more apoptotic and tumour-suppressive signalling.

#### Cold breast tumours show increased expression of immunosuppressive factors

Our data revealed that the luminal tumour co-marker progesterone receptor (PR), which is linked to an immunosuppressive tumour microenvironment [47, 48], displayed a decrease in expression at the highest significance as compared to the cold subgroup (Figure 4B, C). Another contemplated immunomodulation factor is the PPARγ/RXRα pathway which regulates cell proliferation and inflammation [49]. It has been implied that expression of Peroxisome proliferator-activated receptor γ (PPARγ), a key modulator of this pathway, correlates with supressed immunosurveillance [50-52]. Interestingly, a positive correlation was found for phosphorylated PPARγ (pS112) and PR (Figure 5B, Pearson’s r=0.45, P<0.0001), indicating a co-expression of both factors by cold carcinomas.

**Figure 5.**
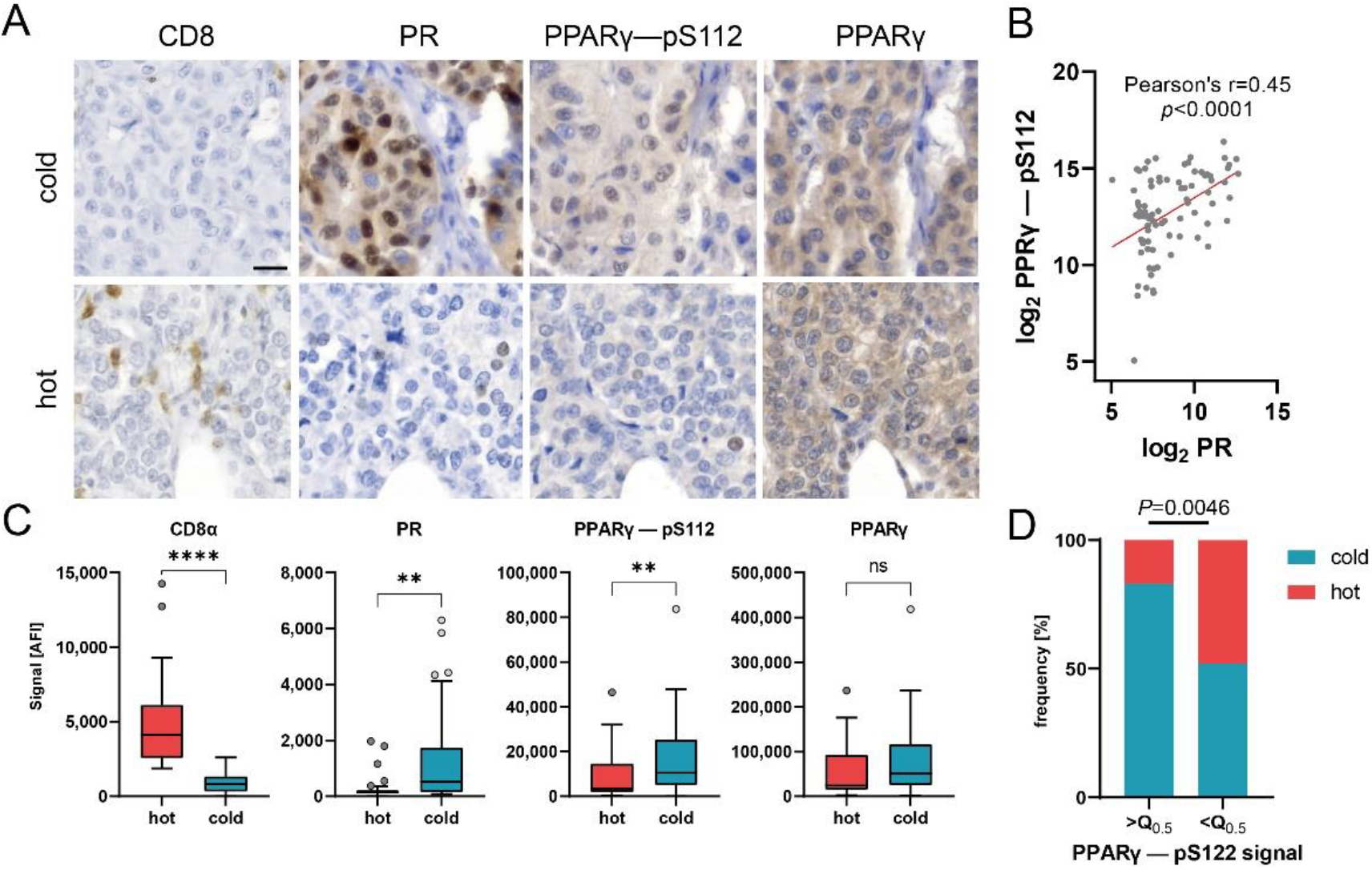
PPARγ phosphorylation correlates with PR expression and may impair immune infiltration. **(A)** Representative images of CD8, PR, PPARγ-pS112 and PPARγ immune-histochemical staining in hot and cold carcinomas. Scale bar, 20 µm. **(B)** Scatter plot of log2 transformed DigiWest signal for PPARγ-pS112 and PR. Pearson correlation, r=0.45, P<0.0001. **(C)** Box plots showing protein expression for CD8α, PR, PPARγ-pS112 and PPARγ in cold (n=57), hot (n=27) and baseline (n=10) samples. Mann-Witney-*U* test, **P<0.01, ****P<0.0001, ns indicates no significant difference. **(D)** Distribution of hot and cold status in carcinoma samples with PPARγ—pS112 expression higher or lower than median expression. n=42. Q_0.5_ = 7004 AFI. Fishers exact test, P=0.0046.

Additionally, our results show that expression of the phosphorylated variant of PPARγ (pS112), but not the total protein variant, was significantly upregulated in cold tumour samples, whereas CD8 expression displayed an opposing trend (Figure 4C, Figure 5C). Immuno-histochemical staining of representative samples verified the predominant expression of PPARγ (pS112) and PR by cancer cells in cold tumour samples (Figure 5A). Concomitantly, the percentage of samples classified as cold tumours was found to be significantly enriched in samples with an elevated (greater than median) PPARγ-pS112 signal (Figure 5D, Fisher’s exact test, P=0.0046). Hence, our data suggest that PPARγ phosphorylation might be involved in a mechanism governing immune cell repulsion in breast cancer.

## Discussion

Mutational changes in cellular signalling that are triggering cell growth are key events in the transformation process that lead to the formation of tumour cells. Targeted analysis of central signalling pathways helps to track the effects of such mutations and can be used to functionally classify tumours by subtype. The DigiWest methodology employed here is capable of achieving this on a much broader scope compared to classical immune-histological approaches and the knowledge generated by characterising the activity of central signalling pathways can be utilised to identify novel targets for therapeutic intervention [53].

Here we used a well-characterized collection of archived breast cancer tissues for targeted protein profiling and aimed to concomitantly detect (i) signalling proteins and their activated variants and (ii) immune cell markers that define the tumour microenvironment. Thereby, we were able to screen for aberrations in the intra- and extracellular communication and to assign an immune status of each individual tumour tissue. Since patient specific follow-up data for all samples was available and could be integrated into the analysis the correlation between protein expression levels, immune status and tumour reoccurrence could be analysed.

Immune cell infiltration of breast cancer tissue and stroma is linked to a better prognosis [6]. On the contrary, non-immune infiltrated tumours show no or low response to current immune therapy [54]. To survey for the presence of infiltrating immune cells in cancer tissue, we measured central immune cell markers (CD8a, CD11c, CD16, and CD68) simultaneously, and categorized the present cohort in highly-infiltrated (“hot”) tumour and lowly-infiltrated (“cold”) tumour samples. Assessment of patient outcome data revealed a significant difference in event-free survival in favour of the highly immune cell infiltrated group. Significantly, higher amounts of PD-1 and additional immune cell markers were detected in these samples indicating a higher immunosurveillance. Conversely, the specific Treg cell marker FoxP3, necessary for the immune-suppressive activity and immunological tolerance [55-57] was found to be enriched in cold breast tumours. This indicates that in this cohort, higher expression of FoxP3+ cells leads to retention of immune cell infiltration within tumour tissue and consequently poorer patient outcome.

At the same time, we could show that highly immune cell infiltrated tumour samples display elevated immunological signalling activity, as was indicated by upregulation of regulatory proteins Jak2, STAT4 and STAT1 including its activating phosphorylation at Tyrosin 701. These crucial members of the Jak/Stat pathway are important for cytokine response and constitute key regulators of the immune system [58, 59].

In a recent study, high apoptotic activity in breast cancer tissue was shown to be associated with high infiltration of immune cells. Based on mRNA expression data the authors hypothesize that increased apoptosis is associated with immune cell killing [60]. Here we report that in hot breast cancer tissue, central apoptotic marker proteins like the cleaved version of initiator Caspase 9, the cleaved effector Caspase 6 and Bax were upregulated. Increased levels of p53 phosphorylated at Ser20 and Histone H3 phosphorylated at Ser10 in these tumours indicate a likely involvement of DNA damage [40].

Phosphorylated peroxisome proliferator-activated receptor (PPARγ-pS112) was present in higher amounts in samples with lower immune cell infiltration. Besides its common function in adipogenesis and lipid metabolism [61], PPARv has been linked to evasion of immunosurveillance and impairment of CD8 T-cell infiltration in muscle-invasive bladder cancer [62]. Therefore, we hypothesize that PPARγ phosphorylation may be involved in an immunosurveillance evasion mechanism employed by breast cancer cells. Hence, PPARγ may serve as a potential marker for patient stratification or as a target for therapeutic intervention. Yet, additional research is required to elucidate this question further.

The application of the DigiWest technology enabled us to review the receptor status, immune cell infiltration and protein expression of approx. 150 proteins and protein variants in parallel from minimal amounts of fresh frozen breast cancer biopsies, demonstrating the unique potential held by this approach.

## Ethics approvals

The use of retrospective human samples is ethically approved (788/2018BO2) by the ethics commission at the Medical Faculty of Tuebingen University.

## Informed Consent Statement

Informed consent was obtained from all subjects involved in the study.

## Author contribution

Conceptualization, F.R., A.K and M.T; methodology, F.R., and M.T; formal analysis F.R, N.K and M.T; establishment of original pathological diagnosis and receptor profile, review of the adequacy of tumour cell content, A.S.; investigation, F.R., S.J and N.A; data curation F.R., N.K, M.T; writing—original draft preparation F.R., Aa.S, N.K. and M.T.; writing—review and editing, C.S., A.H., M.H., K.S.-L; A.K. provided material support; supervision, M.T., S.Y.B. and K.S.-L; project administration, C.S. All authors have read and agreed to the published version of the manuscript.

## Data availability statement

The data that support the findings of this study are available from the corresponding authors upon reasonable request.

## Supporting Material

**Supplementary Figure S1.**
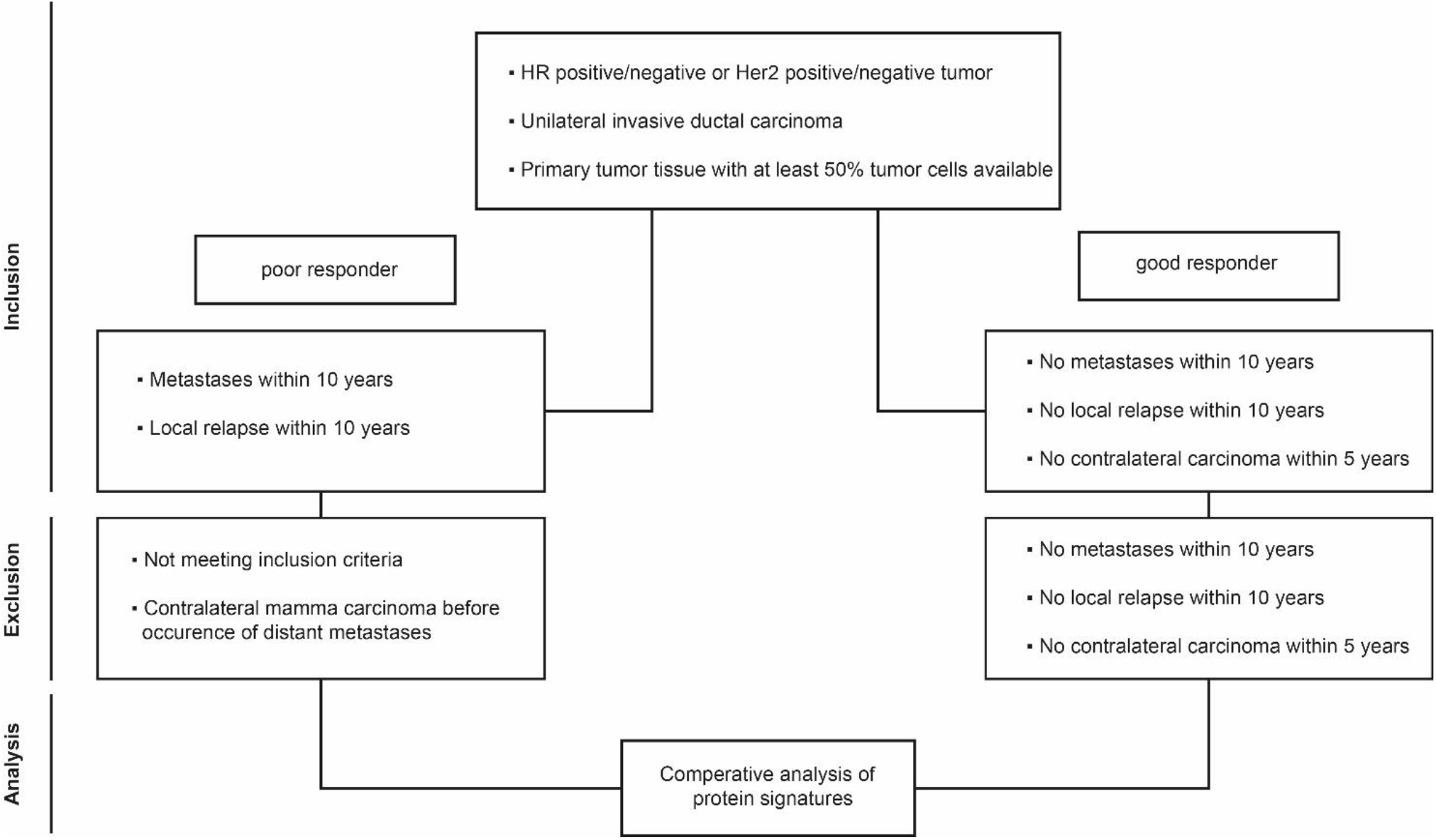
Study flowchart.

**Supplementary Figure S2.**
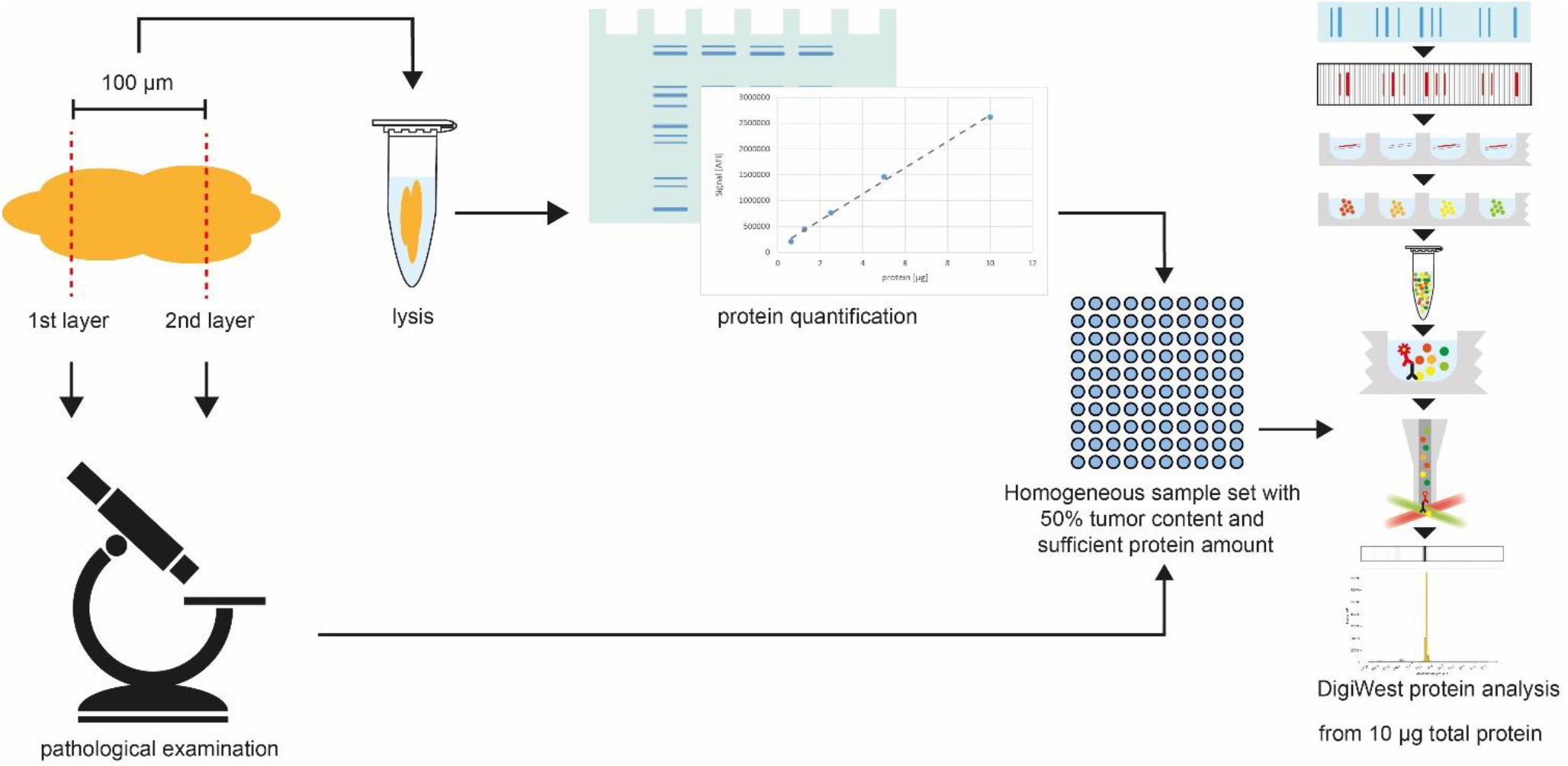
Schematic depiction of study workflow. Layered cuts of tumour biopsies were generated, and intermediate tissue (100 µm) was collected. A pathologist assessed tumour content in first and second layer. Collected sample material was lysed and protein quantification was performed. Samples with tumor content of ≥50 % tumorcontent and sufficient protein amount were selected for DigiWest analysis.

**Supplementary Figure S3.**
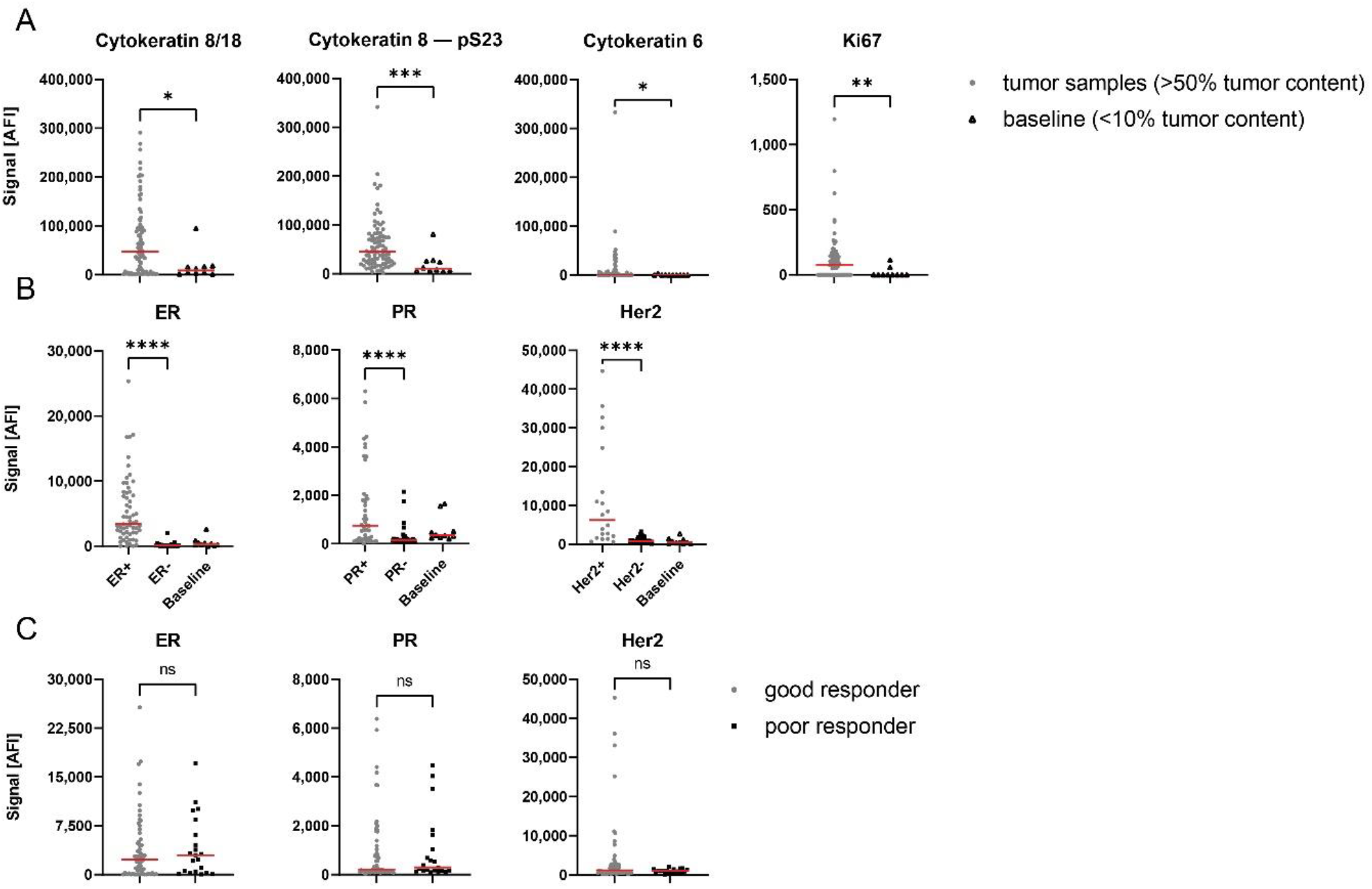
Tumour marker and receptor expression in baseline samples versus tumour samples. **(A)** Cytokeratin 8/18, Cytokeratin 8 – pS23, Cytokeratin 6 and Ki67 expression as scatter plots in samples with ≥ 50 % tumour content (Tumor samples, n=84) and ≤10% tumour content (baseline, n=10). **(B)** Scatter plots showing protein expression of ER, PR and Her2 in respective receptor-positive or negative and baseline subgroup (ER+ n=60; ER-n=24; PR+ n=46; PR-n=38; Her2+ n=20; Her2-n=63) as well as **(C)** in good (n=58) and poor responders (n=21). In **A, B, C** Mann-Whitney-*U* test, ****P<0.0001; ***P<0.001; **P<0.01; *P<0.05; ns indicates no significant difference.

**Supplementary Figure S4.**
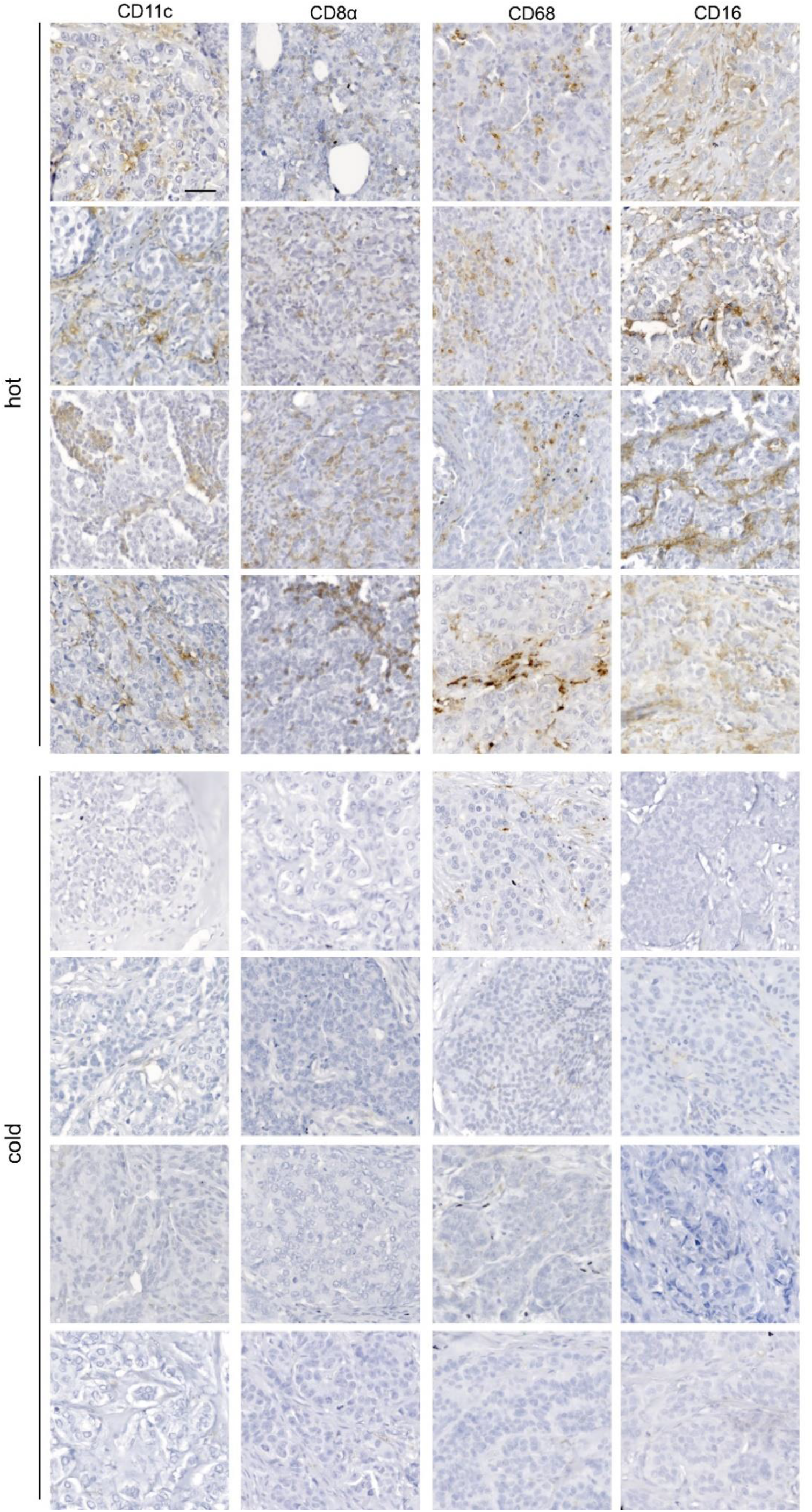
Overview of IHC staining of immune cells. Representative images of CD11c, CD8, CD68 and CD16 immuno-histochemical staining in hot and cold samples. n=4. Scale bar, 50 µm.

**Supplementary Figure S5.**
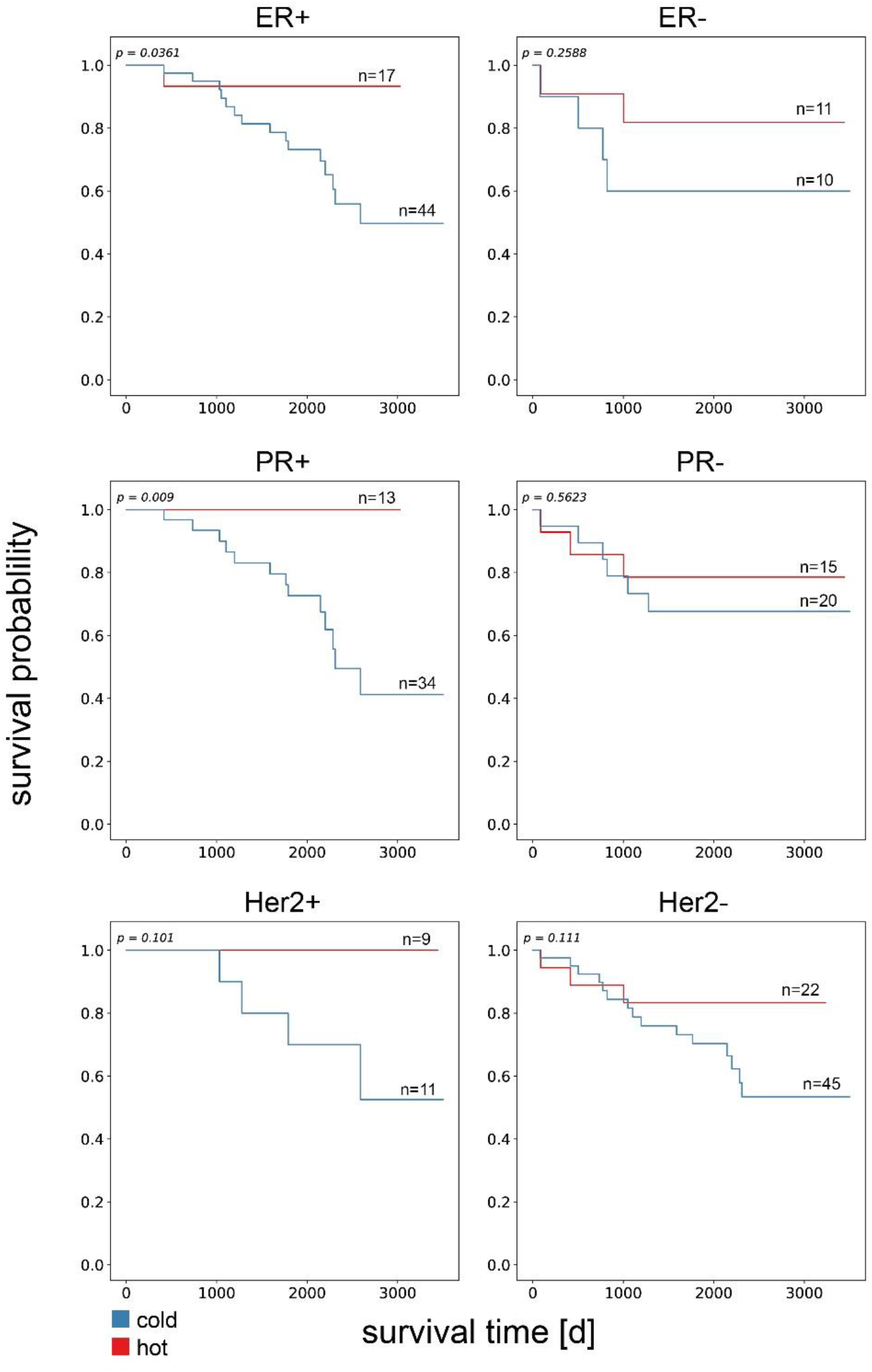
Influence of infiltrating immune cells on event-free survival in hormone receptor and Her2 positive/negative tumours. Kaplan-Meier analysis of event-free survival between hot and cold carcinoma samples in ER, PR and Her2 positive and negative subgroups. A significant difference in EFS was found in ER+ and PR+ subgroup (P<0.05). Log-rank test.

**Supplementary Figure S6.**
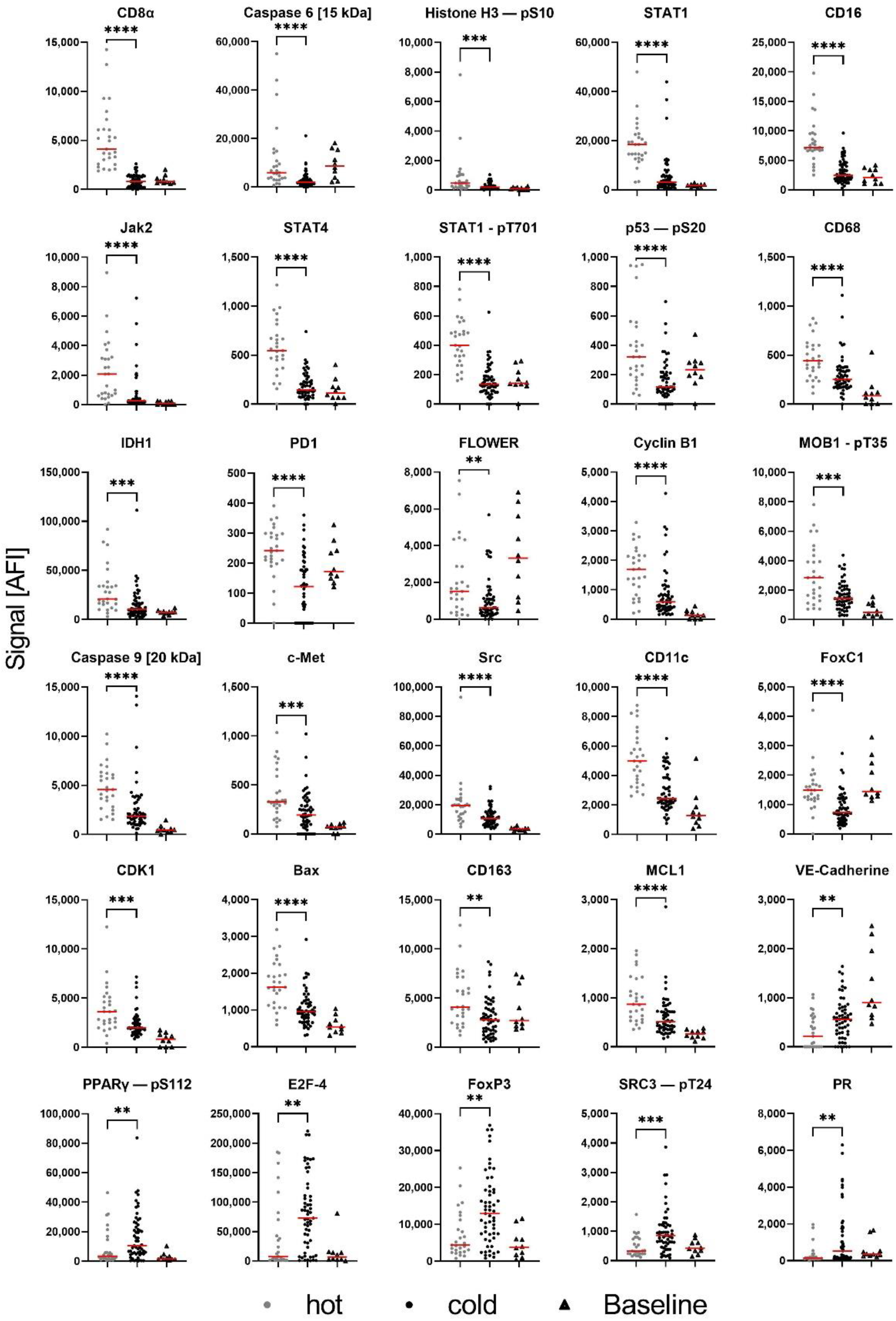
Differences in protein expression between hot and cold tumors. Protein expression of 30 analytes for hot (n=27), cold (n=57) and baseline (n=10) subgroup which revealed significant differences in protein expression and a fold change of at least 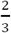 between hot and cold samples. Mann-Whitney-*U* test, ****P<0.0001; ***P<0.001; **P<0.01.

**Table S1.** Correlation values for all measured immune cell markers. Spearman’s correlation.

|  | **CD4** | **CD25** | **CD56** | **CD163** | **CD11c** | **CD68** | **CD16** | **CD8a** |
| --- | --- | --- | --- | --- | --- | --- | --- | --- |
| **CD4** | 1 | -0.2 | 0.4 | -0.4 | 0.1 | 0.2 | 0 | 0 |
| **CD25** | -0.2 | 1 | 0.4 | 0.2 | 0.1 | 0.2 | 0.5 | 0.5 |
| **CD56** | 0.4 | 0.4 | 1 | 0 | 0.4 | 0.5 | 0.4 | 0.5 |
| **CD163** | -0.4 | 0.2 | 0 | 1 | 0.5 | 0.4 | 0.3 | 0.4 |
| **CD11c** | 0.1 | 0.1 | 0.4 | 0.5 | 1 | 0.7 | 0.6 | 0.6 |
| **CD68** | 0.2 | 0.2 | 0.5 | 0.4 | 0.7 | 1 | 0.7 | 0.6 |
| **CD16** | 0 | 0.5 | 0.4 | 0.3 | 0.6 | 0.7 | 1 | 0.7 |
| **CD8a** | 0 | 0.5 | 0.5 | 0.4 | 0.6 | 0.6 | 0.7 | 1 |

